# Xi rhythms: decoding neural oscillations to create full-brain high-resolution spectra parametric mapping

**DOI:** 10.1101/2019.12.17.880328

**Authors:** Shiang Hu, Pedro A. Valdes-Sosa

## Abstract

Neural oscillations excitability shape sensory, motor, and cognitive processes of the brain operation. To quantify the spectrum of brain magnetic and electrical recordings has become the main methodology to study neural oscillations. However, there is still lacking a valid approach, although the literatures on the spectrum decomposition is vast. In this work, we fit the neural spectrum by means of the Expectation Maximization algorithm, where the E step turns into a Winner filtering to separate multiple components and the M step is to fit each component by minimizing the smoothness penalized Whittle likelihood and shape-restricted regression, say, monotonicity, that is able to fit the diverse shape of each component. The decomposition allows characterizing the oscillation amplitude, resonance frequency, bandwidth, skewness, kurtosis, and slope of each component. This approach is termed as ‘Xi rhythms’, with Xi and rhythms standing for the background activity and multiple peaks. We apply it to: 1) multinational EEG database consisting of 535 subjects to create the quantitative spectrum norms (QSN); 2) large sample intracranial EEG (iEEG) dataset to infer the oscillations from recorded region to unrecorded areas within one subject and over inter-individuals and create the full brain high resolution statistical spectra parametric mapping. The statistical spectrum parameter mapping of iEEG promisingly provides an atlas and creates a norm for neural oscillations and quantitative electrophysiology study which can gain us more insightful understanding to brain dynamics, cognitive process and mental disorders.

## 1. Introduction

Neural oscillation shapes the extracellular fields and currents accumulated by the synchronous neural mass activities and is recorded passively by the Electroencephalography (EEG), intracranial EEG (iEEG), Electrocorticography (ECOG), and Magnetoencephalography (MEG) (Buzsáki et al., 2012). The nature of cognitive process, such as attention, memory, consciousness, the generation of brain dysfunction, mental disorder, pathophysiology, e.g. schizophrenia, and the cortical networks are revealed by the study to neural oscillations (Buzsaki, 2004; Uhlhaas and Singer, 2006, 2010; Ward, 2003). The spectrum analysis of magnetic-electric recordings provides the fingerprints of large scale neural oscillations (Siegel et al., 2012).

Spectrum decomposition and building the model to characterize the brain rhythms have been a hot issue since 1970s, when the conception of quantitative EEG was proposed (John et al., 1977; Lopes da Silva et al., 1974; Zetterberg, 1969). Zetterberg mainly employed the idea of ‘filtering’, recognized three types of components in the spectrum, and estimated the parameters with maximum likelihood estimate (Isaksson et al., 1981; Wennberg and Zetterberg, 1971; Zetterberg and Ahlin, 1975). With the need of computer assisted differential diagnosis in quantitative EEG (John et al., 1988), Pascual and Pedro Valdes etc. put forward a likelihood ratio test based approach named as ‘Xi alpha’ model to extract the oscillation parameters for the background activity --- the ‘Xi’ process and the alpha rhythm, and extended to multichannel spectra analysis (Pascual-marqui et al., 1988). Xi alpha model used student t curve to fit each component and only can fit the alpha peak. Recently, FOOOF is developed to the parametrize the 1/f process and the multiple peaks with gaussian kernels in a heuristic Backfitting approach by minimizing the fitting residuals of sum square (Haller et al., 2018).

However, the drawbacks of the existing studies could be rethought from the following: 1) it is still not clear that whether the background oscillation can be described 1/f process (Abry et al., 1995); 2) due to the large diversity of multiple peaks (rhythms) in shape, is the parametric fitting such as previously adopted student t curve and gaussian kernel effective/robust to fit all the peaks? 3) is there other theoretical valid criterion to evaluate the goodness of spectrum estimation than least square used in FOOOF? 4) how many rhythms (peaks) are ongoing in a certain cognitive process, say, how many peaks should one fit given the estimated spectrum?

To address the problem of single component fitting on its diverse shape, smoothed unimodal regression seems to be a promising way due to its non-parametrization and flexible restriction on shape (Eilers, 2005; Pya and Wood, 2015). Considering the shape properties of spectrum components, the background oscillation is just a monotonically decreasing function and the peaks are first monotonically increasing to the peak location and then monotonically decreasing function. To seek for a valid criteria of spectrum estimation, we found that Whittle likelihood is a consistent estimator with the assumption that the Fourier coefficients at a specific frequency bin is circularly complex normal distributed given the Central limit theorem (Whittle, 1953; Whittle et al., 1951). One of the keys to the spectrum fitting is the number of peaks, we considered generally two ways: i) data driven, the number of peaks is just the size occurred in the spectrum curve; the very close double peaks with one minor trough in the middle is considered double peaks; in this case, best fitting is obtained with maximum likelihood estimate or least fitting residuals of sum square ii) model driven, the number of peaks doesn’t depend on its size occurred; the neighboring peaks with one minor trough may be taken as one peak; this relies on more priors from the studies of neural oscillations; in this case, either overfitting or underfitting should make sense but not only seeking for minimized fitting residuals of sum square; it is implemented by the Expectation-Maximization (EM) method with group lasso incorporating the neurophysiological priors.

Up to now, only the fitting and the decomposition has been described. Whereas, decoding (quantifying) neural oscillations (spectrum) should be taken in the systematic view displayed in the figure 1. Fitting and decomposition breaks the neural spectrum into pieces of spectrum components which is in fact an inverse problem where EM algorithm (a particular case of majorization-minimization (MM)) is advantageous (Demidenko, 2005; McLachlan and Krishnan, 2008; Zhou et al., 2015). Some parameters from the spectrum components are extracted as interpretable indicators of neural oscillation, such as the oscillation amplitude, resonance frequency, bandwidth, skewness, kurtosis, slope, and relative power, say the prominence to the total power. This will allows the application of Xi Rhythms into 1) building oscillation norms over sensor and source space with large normative database; 2) prediction and differential diagnosing as biomarkers for cognitive process and mental order; 3) spatial inference with iEEG/ECOG data from recorded region to unobserved areas over the intrasubject level and the inter individual subjects level as well (Owen and Manning, 2017); 4) the full brain high resolution statistical spectrum parametric mapping may provide a norm prior for the brain magnetic-electrical inverse source imaging.

**Figure 1.**
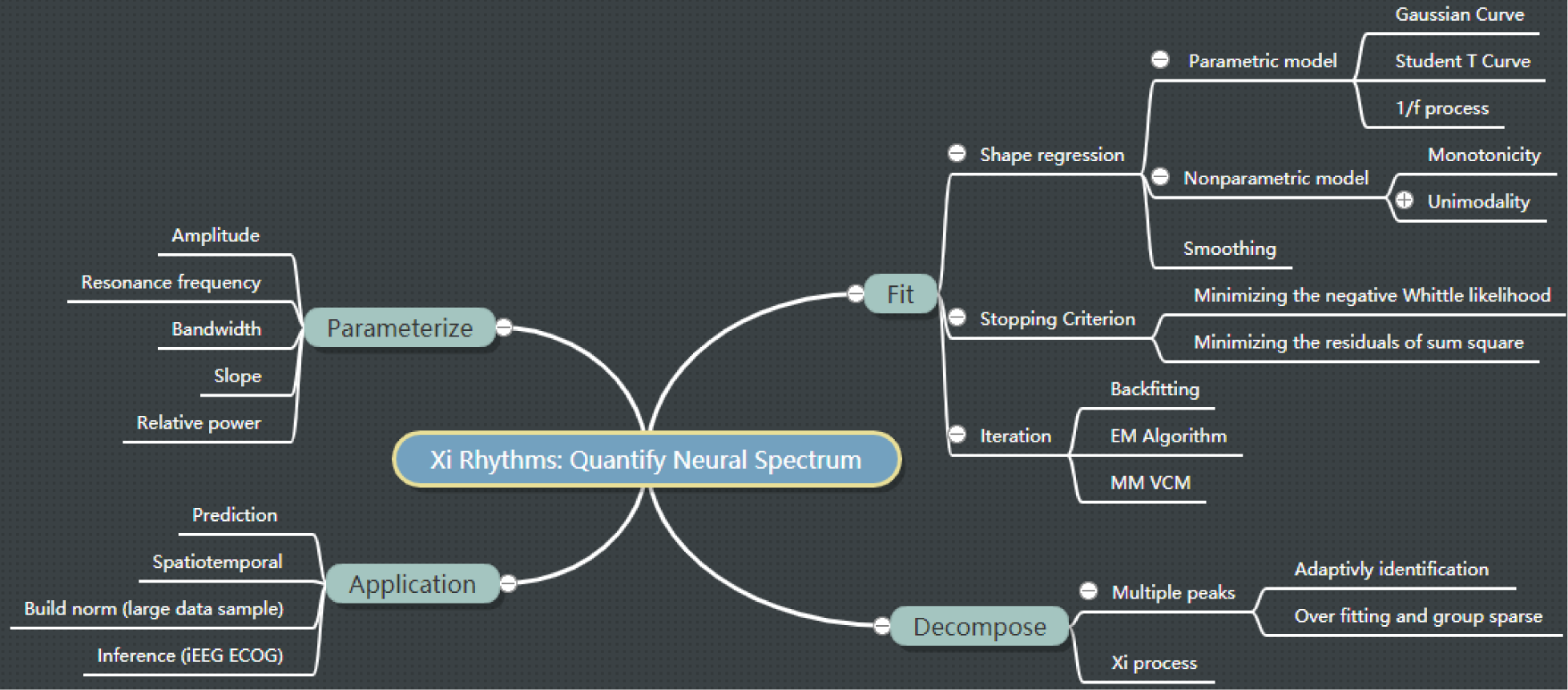
A systematic view of Xi Rhythms on quantifying the neural spectrum.

In this paper, we show that the E step in spectrum fitting acts as a Winner filer to separate the spectrum components and the M step is to minimize the shape constrained and smoothness restricted Whittle likelihood. A multinational quantitative oscillation norm is built with the collected EEG data from Cuba, United states, Switzerland and Mexico. A full brain high resolution spectrum parametric mapping (SPM) is created thanks to the shared iEEG dataset (Frauscher et al., 2018; Owen and Manning, 2017).

## 2. Methods

In what follows, we denote scalars with lowercase symbols (e.g. *x*), vectors with lowercase bold (e.g. **x**), matrices with uppercase bold (e.g. **X**); unknown parameters will be denoted by Greek letters (e.g. *ξ*). Furthermore, **1** is the vector of ones; 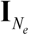 is a *N*_*e*_ by *N*_*e*_ identity matrix; *N*^*C*^(**μ**, **Σ**) is the circularly complex multivariable Gaussian distribution with mean vector **μ** and covariance matrix **Σ**; (⋅)^**T**^ is the transpose of (⋅); **X**^+^ is the pseudo-inverse of **X**; *tr*(⋅) is the trace of (⋅); 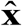 is the estimation of **x**; ||⋅||_2_, ||⋅||_*F*_, and ||⋅||_*M*_ are the Euclidean norm, the Frobenius norm, and the Mahalanobis norm, respectively; *E*(⋅) is the expectation operator; ⊙ refers to the Hadamard product.

### 2.1. Decomposition

In magnetic-electrical recordings spectrum analysis, the recordings are segmented, and each segment is taken the discrete Fourier transform. If *y*_*ω*_ is the Fourier coefficient at the frequency *ω* of the data at segment *s* (*ω* = 1,…, *n*_*ω*_ and *s* = 1,…, *n*_*s*_), **z** is a *n*_*k*_ ×1 vector of ones, indicating the linearly additional effect of *n*_*k*_ components, and **b**_*ω*_ is a *n*_*k*_ ×1 vector, consisting of the Fourier coefficients of *n*_*k*_ components at frequency *ω* and epoch *s*, 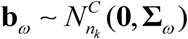, **Σ**_*ω*_ = *diag*(**σ**_*ω*_), 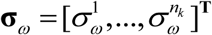, *ε ~ N*^*C*^(0, *σ*_*ε*_), is the decomposition error with a constant variance across frequencies, then the Fourier coefficient **decomposition model** is

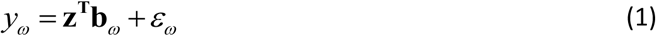

And estimated spectrum is the sample variance over segments,

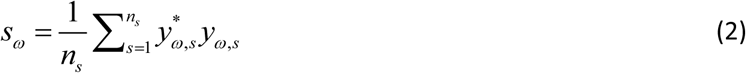

Given the real spectrum as *s*_*ω*_ and the theoretically fitted spectrum as *σ*_*ω*_, the **Whittle likelihood** is define as (Pawitan et al., 1994; Whittle, 1953; Whittle et al., 1951)

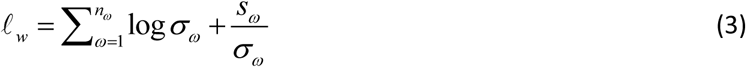

The spectrum decomposition problem transforms into the estimation of covariance of **b**_*ω*_ given the observation model (1). By means of the variance components model and Expectation-Maximization algorithm (McLachlan and Krishnan, 2008), the vector of unknow parameters as 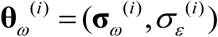, the complete the negative log likelihood is

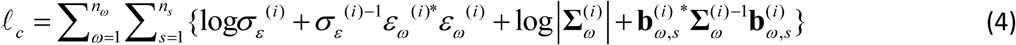

#### E step

The variance of *y*_*ω*_ in (1) is 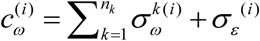. Taking the minimum norm least square solution and matrix inversion lemma (Tarantola, 1987), we obtain

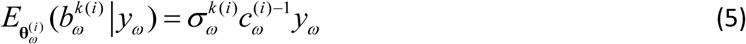

which apparently shows that the spectrum decomposition is a Winner filtering problem.

In the E step, it is also derived

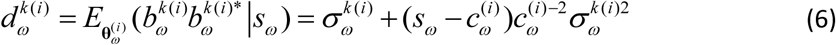

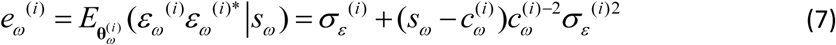

#### M step

The conditional complete negative log likelihood is

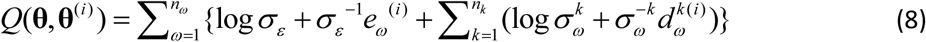

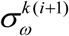 is an approximated estimator to 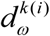 and fitted by the smoothness constrained shape restricted Whittle likelihood, which is described detailly in the next section.

Assuming that the variance for the Fourier coefficient error in (1) is constant for all the frequencies, it is calculated in the M step as

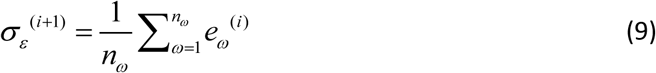

#### Incomplete loglikelihood

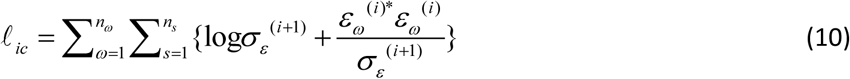

The EM algorithm will finish the iterations till the incomplete negative log likelihood converges.

### 2.2. Fitting

The fitting to the 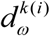 in the M step in the decomposition part is singled out as this section. If rearranging the scalars at each frequency into vectors, we have 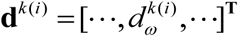 and 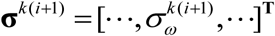. Further, the Whittle likelihood in (3) is expressed as

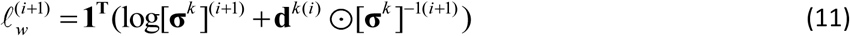

To estimate **σ**^*k*^ in a nonparametric way with smoothness constraint and shape restriction (Eilers, 2005; Wahba, 1980), the objective function is

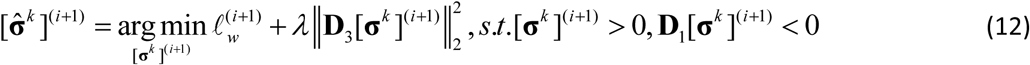

where *λ* is the smoothing parameter, **D**_3_ is the smoothness operator (3^rd^ order differential matrix), and **D**_1_ is the modified 1^st^ order differential matrix. If **d**^*k*(*i*)^ is a motorically decreasing curve, then **D**_1_ is a gradient matrix which keeps the full 1^st^ differential matrix; otherwise for peaks, the sign of the entries on the left side of **D**_1_ is inverted to calculate the negative gradient and positive gradient around the maxima, say, the peak maxima splits the **D**_1_ with negative gradient operator in the left side of the peak.

#### Smoothing parameter selection

Selecting a proper smoothing parameter is essential to fit a smooth curve to each component.

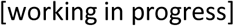

### 2.3. Characterizing

#### 2.3.1 Oscillation amplitude (OA), resonance frequency (OF), full width half magnitude (OW)

**Figure 2.**
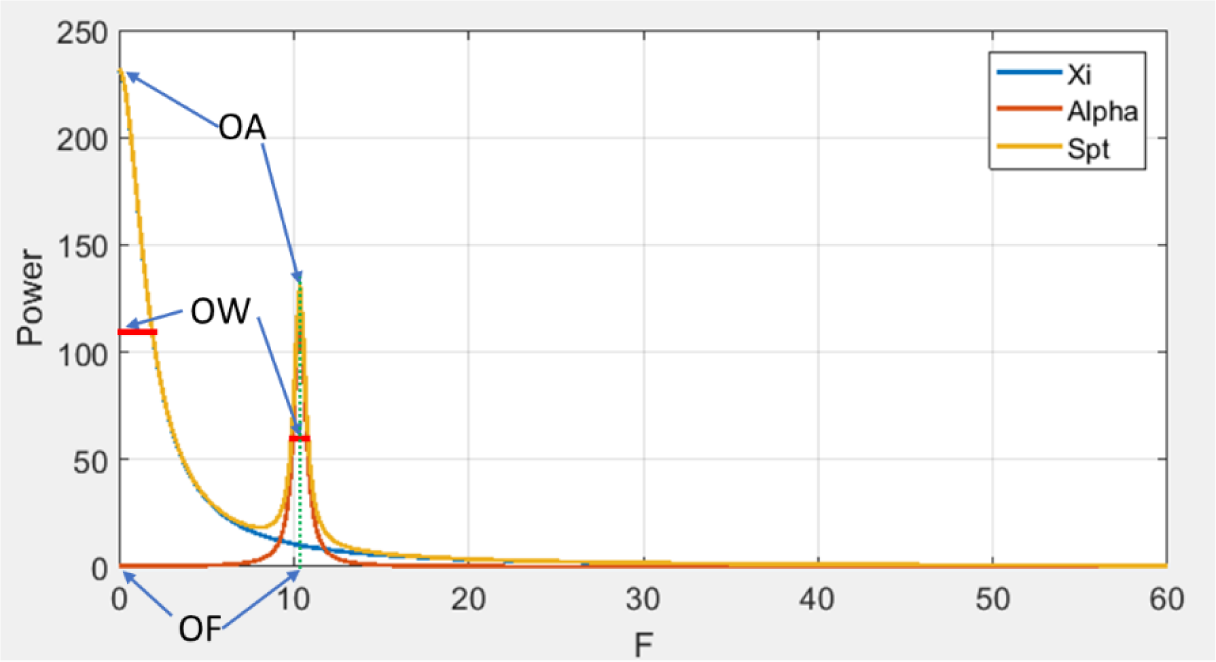
an example to capture the oscillation amplitude, resonance frequency, width.

#### 2.3.2 Oscillation skewness (OSK), slope (OSL), and kurtosis (OKT)

OSK is a measure of asymmetricity of peaks. OSL measures the sharpness of a peak. OKT is a measure of the tail of a peak.

**Figure 3.**
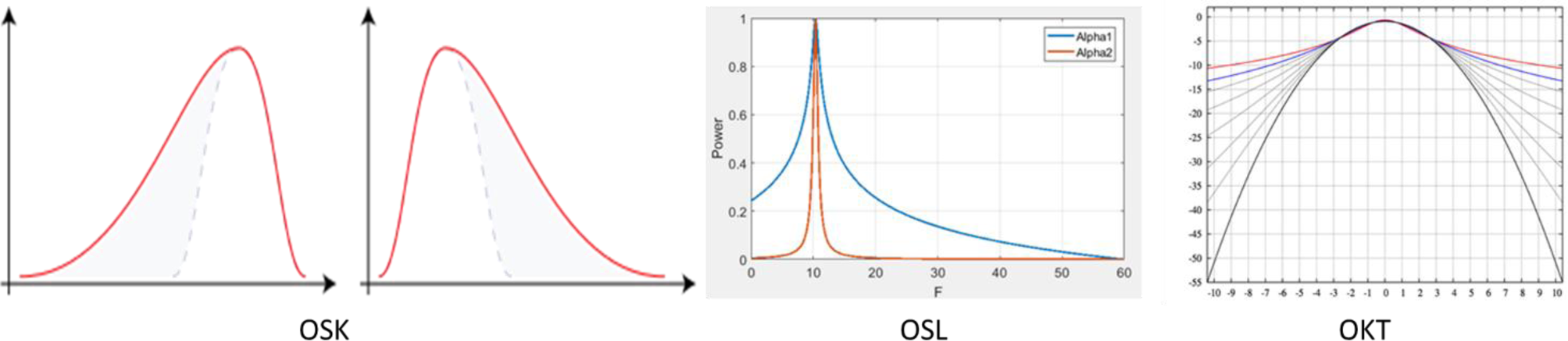
Oscillation skewness (OSK), slope (OSL), and kurtosis (OKT)

### 2.4. Extending to cross-spectra analysis

Xi Rhythms is extended to multichannel cross-spectra analysis. The multivariate Xi Rhythms model is expressed as

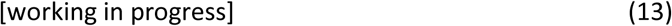

This will allow us to analyze the coherence matrix and build spectra parameters-based connectivity.

### 2.5. Full Brain Spectra Parametric Mapping

The data we analyzed are the open intracranial EEG dataset (Frauscher et al., 2018). Spectra parameters are extracted for each component. We build the graphic model for the ‘inflated’ mesh. Then, all the parameters from the recorded region can be extrapolated to the unrecorded areas. Meanwhile, smoothness over all the voxels are considered.

## 3. Results

### 3.1. Xi Rhythms vs. FOOOF

Xi Rhythms utilizes the signal processing toolbox in MATLAB 2017a. It can adaptively identify the peaks and the valleys. Taking an example of Cuban a spectrum of electrode O1 from Cuban Human Mapping project (Hernandez-Gonzalez et al., 2011), we show the localization and marking with Xi Rhythms,

**Figure 4.**
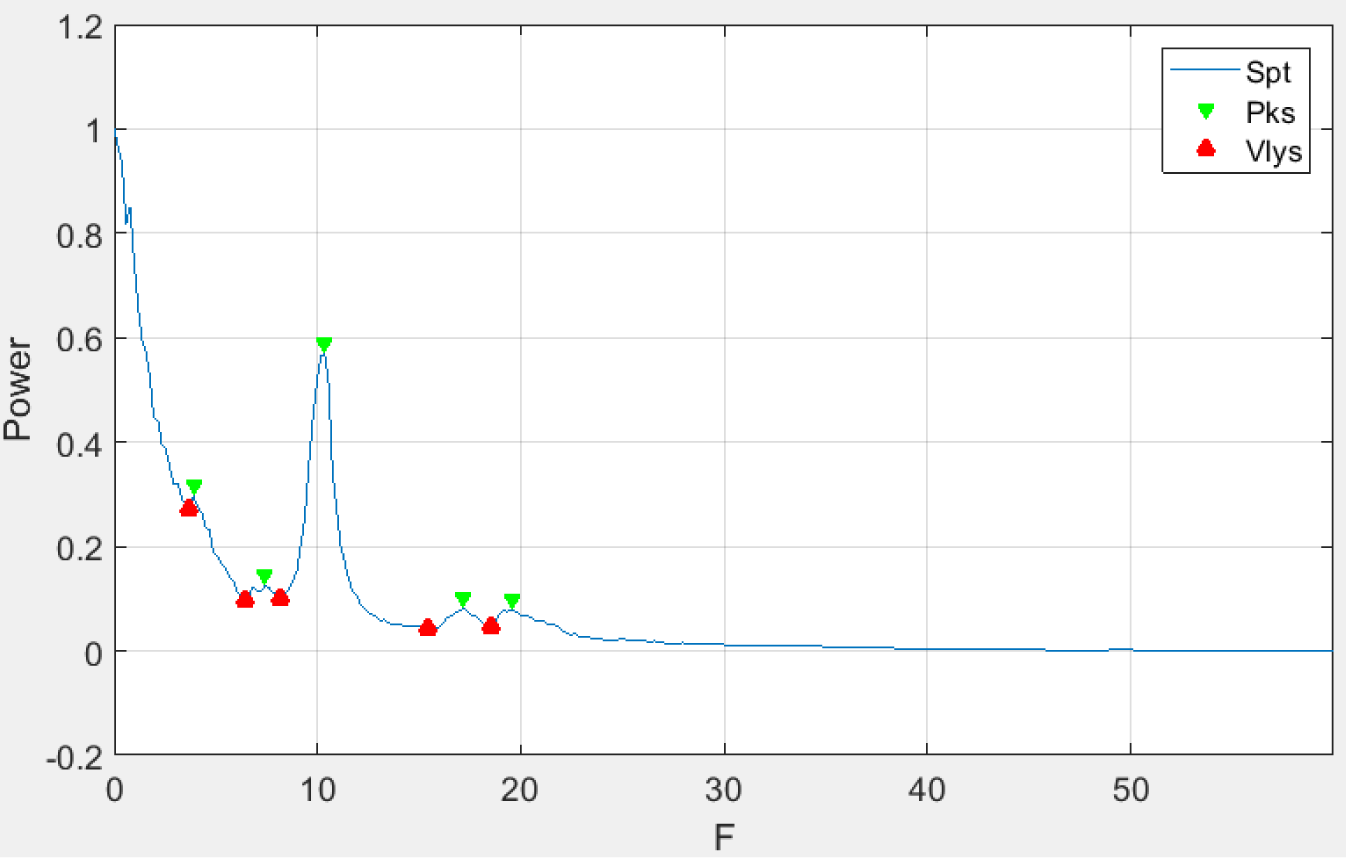
the identification of the peaks (Pks) and valleys (Vlys) with an example

#### Comparison of Xi Rhythms and FOOOF

If input same number of peaks = 3 for both Xi Rhythms and FOOOF, the fitting results is shown in figure 6. We found two visible differences: 1) FOOOF could not cover the alpha peak; however, Xi Rhythms can always cover the alpha peak; 2) with the same number of peaks, FOOOF fits the double peaks as one, whereas Xi Rhythms can keep fitting two peaks.

**Figure 5.**
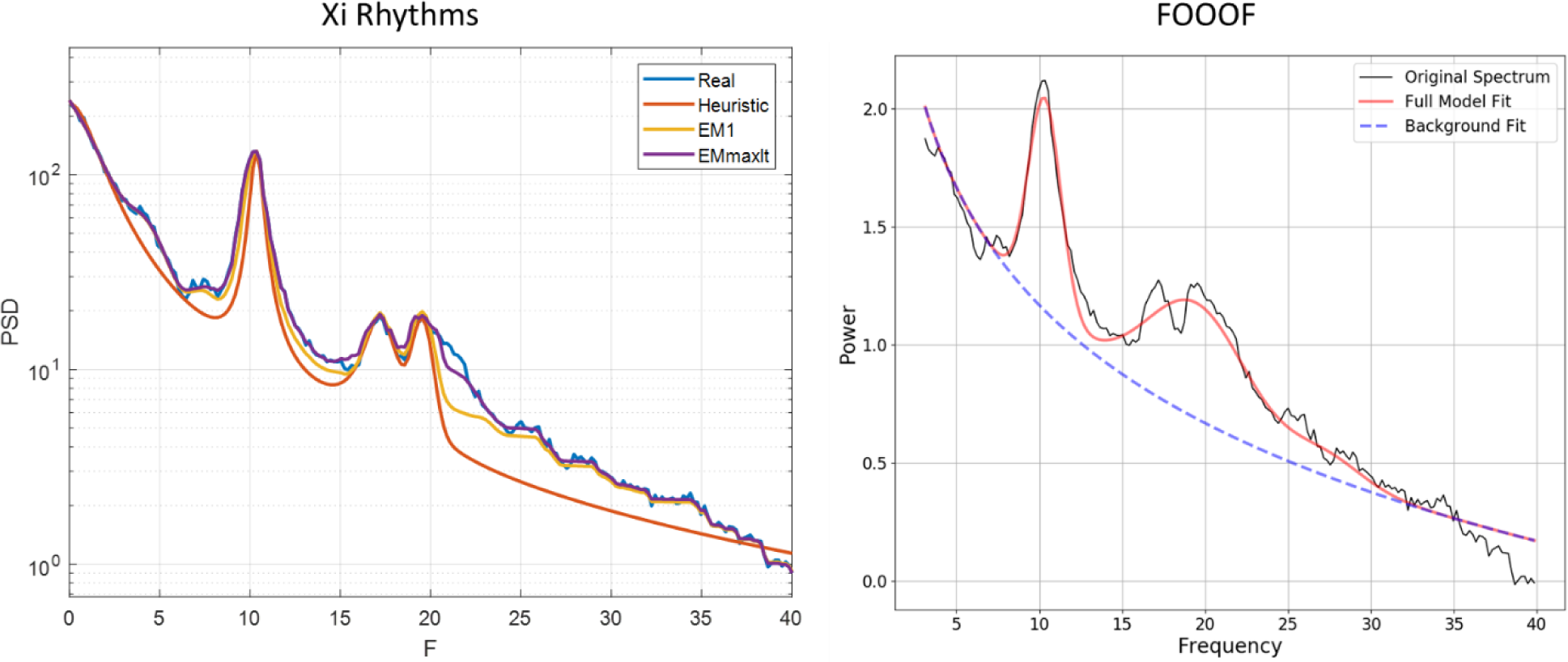
the comparison of Xi Rhythms and FOOOF when fitting the identical real spectrum.

**Figure 6.**
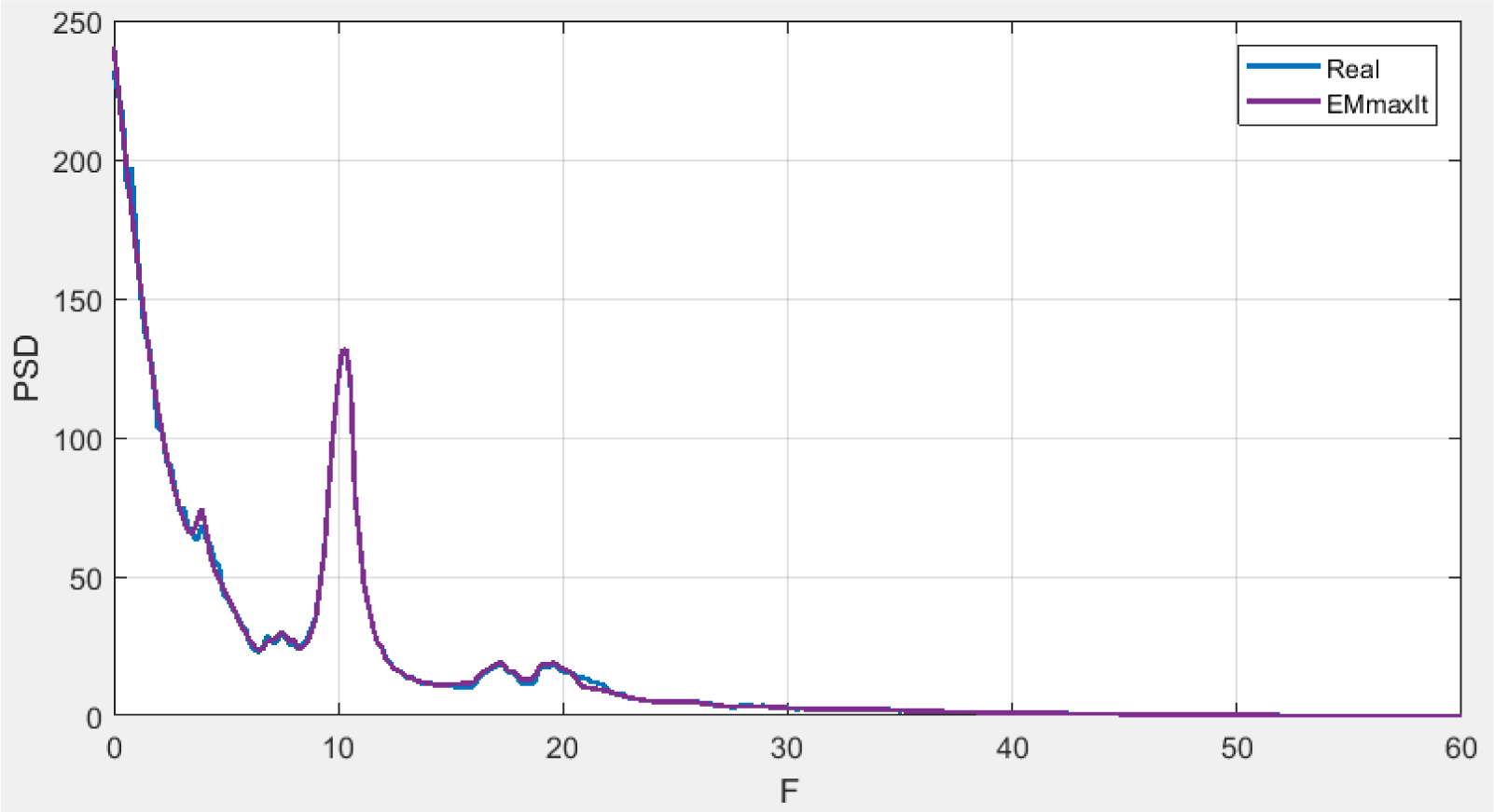
the general fitting to the real spectrum with the same example above

#### Fitting by Xi Rhythms

If inputting number of peaks = 5 which is identified from the figure 4, the general fitting is shown in the figure 6. Apparently, Xi Rhythms can reach a general quite good fitting to the real spectrum. It covers all the peaks and have smoothed background activity.

### 3.2. Multinational quantitative spectrum norms

Multinational dataset is summarized as:

**Figure 7.**
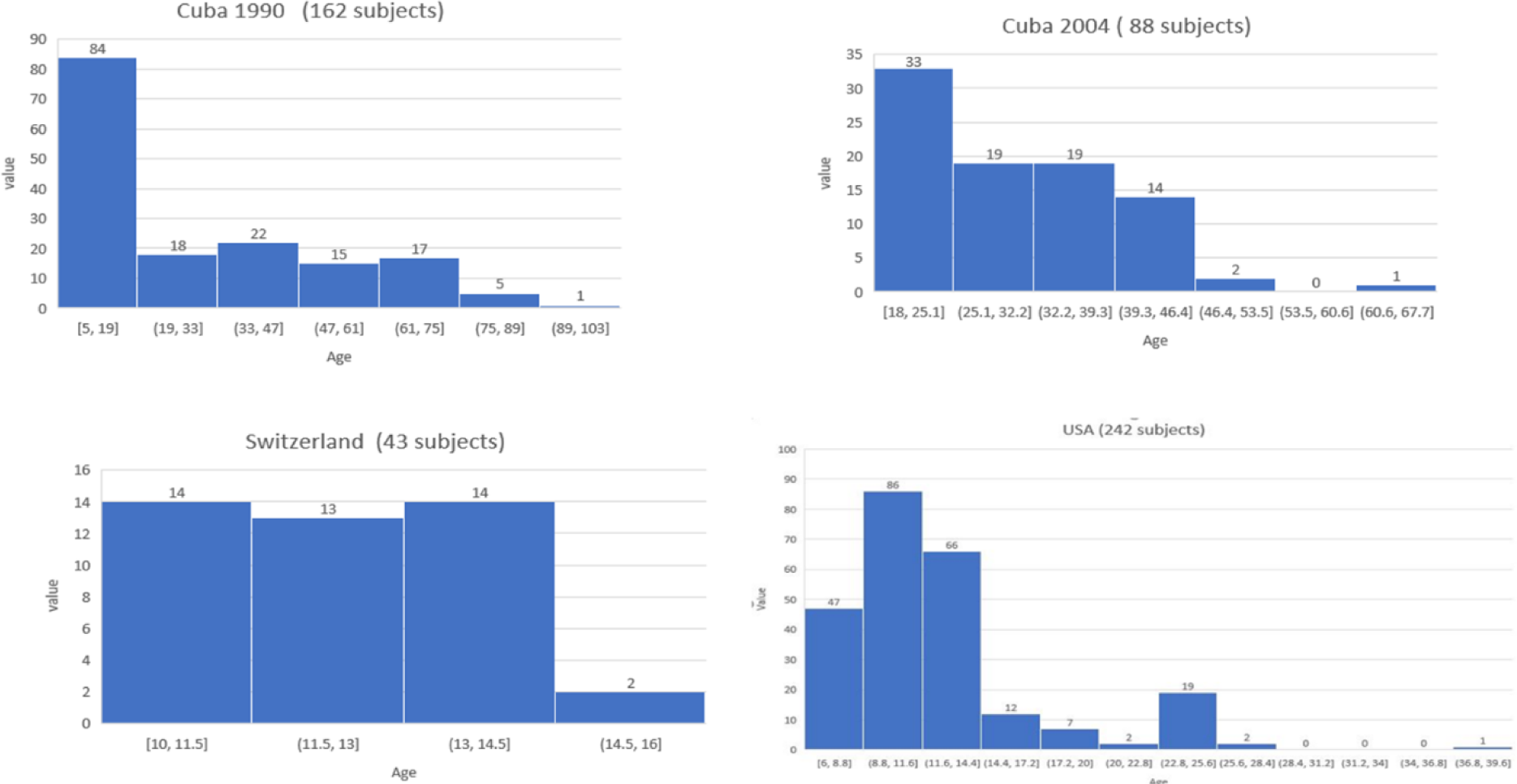
Multinational EEG database summary.

It is conformed that EEG developmental equations are related to age, the stages of brain, and cognitive maturation (Alvarez et al., 1987; Amador et al., 1989; Hudspeth and Pribram, 1990; John et al., 1980). This is reflected by the characteristic of spectra curves, e.g. alpha peaks, are changed over age. Here, the spectrum curves are quantified by the Xi Rhythms, and the extracted parameters are analyzed by forward selection kernel regression method and the mixed effects model (Demidenko, 2005), considering the fixed effects ‘frequencies’, ‘age’, the random effects ‘country’, ‘gender’ and the spectra parameters as response variables. These enables to obtain the refined developmental equations for the spectra parameters.

### 3.3. Full brain high resolution spectrum parametric mapping

We extracted the peak amplitudes over the frequencies (f = 1, …, 40Hz), extrapolated to the full cortical surface. Say, the amplitudes of peaks over all frequencies are fitted to the full brain with smoothness and extrapolation. The result is shown in the figure 8. Prominently, the alpha rhythms peaks localize to the occipital regions, shown under f = 8 - 12 Hz; peaks of delta rhythms (<4 Hz) are located at frontal and occipital regions; the peaks of beta rhythms (13 - 31Hz) appear in the both hemispheres with the symmetrical distribution, and most evidently in the frontal and central partial regions; gamma rhythms (>31Hz) are locating mostly at the left super frontal gyrus and the right lateral super temporal lobe gyrus.

**Figure 8.**
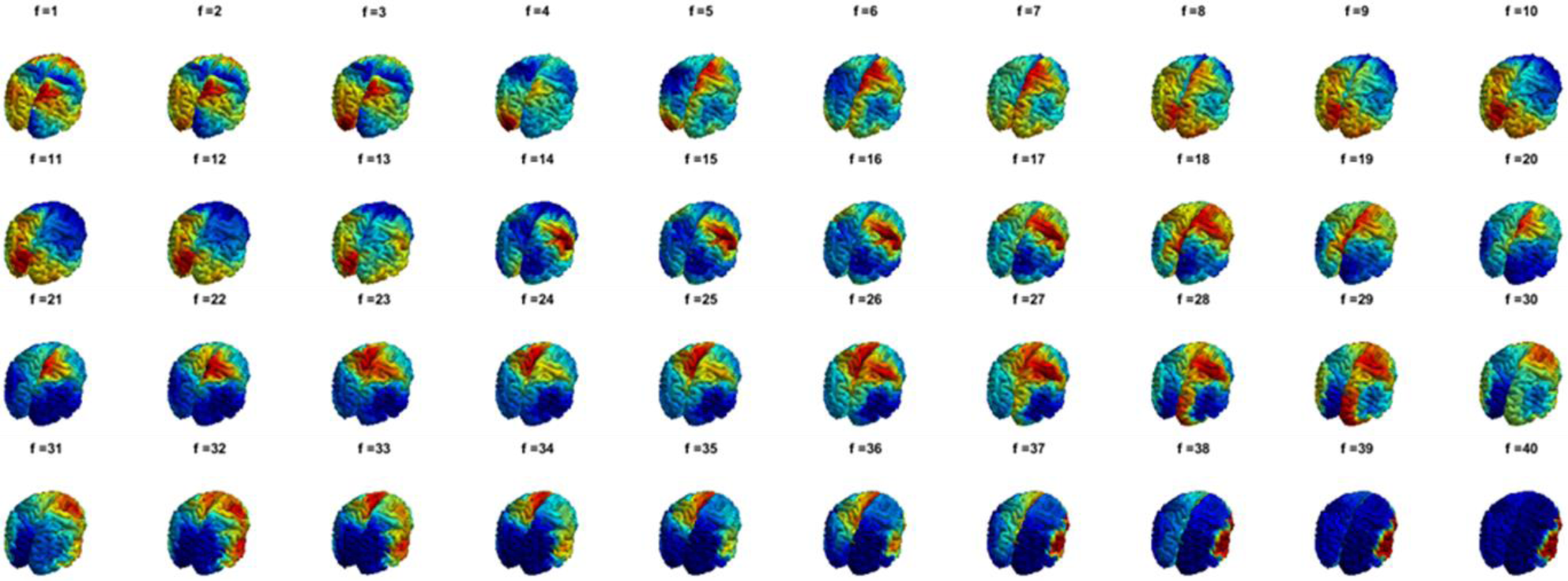
The smoothed peak amplitudes over all the frequencies. The title over each mesh is the frequency in Hz.

## 4. Discussion

There are a few problems remained to solve: (1) Add the constraint **σ** > 0 when using Whittle likelihood at ordinary scale. (2) To tune the smoothing parameter, how to calculate the degree of freedom of a nonparametric fitting? divergence of MLE (Monto Carlo)? (3) Can Whittle likelihood be transformed into a least square, via series expansion or by log transformation (check the expansional family)? (4) Organize the Bayesian and spectrum papers in Mendeley. (5) Review Bayesian analysis, projection estimator, Wahba smoothing and Whittle likelihood.

The developer version of the Xi rhythms toolbox is at https://github.com/ShiangHu/SCMOPT.git.

## 5. Conclusion

Using the EM algorithm and shape regression, Xi Rhythms was proposed to decompose the spectra as the combination of the multiple peaks and the nonperiodic oscillation. The application in the multinational EEG spectral norm and the intracranial EEG dataset illustrated it as an effective neural oscillation decoder to the studies of neural dynamics.

## Acknowledgement

The authors declare no conflicts of interests. This study is supported by the National Science Foundation of China (NSFC), project number: 81601585 and the Canada-Cuba-China project (G0581861128001). We’d like to thank Dr. Thomas Koenig from University for Psychiatry and Psychotherapy Bern, Switzerland for his help on spatial inference.

